# AngularQA: Protein Model Quality Assessment with LSTM Networks

**DOI:** 10.1101/560995

**Authors:** Matthew Conover, Max Staples, Dong Si, Miao Sun, Renzhi Cao

## Abstract

Quality Assessment (QA) plays an important role in protein structure prediction. Traditional protein QA methods suffer from searching databases or comparing with other models for making predictions, which usually fail. We propose a novel protein single-model QA method which is built on a new representation that converts raw atom information into a series of carbon-alpha (Cα) atoms with side-chain information, defined by their dihedral angles and bond lengths to the prior residue. An LSTM network is used to predict the quality by treating each amino acid as a time-step and consider the final value returned by the LSTM cells. To the best of our knowledge, this is the first time anyone has attempted to use an LSTM model on the QA problem; furthermore, we use a new representation which has not been studied for QA. In addition to angles, we make use of sequence properties like secondary structure at each time-step, without using any database. Our model achieves an overall correlation of 0.651 on the CASP12 testing dataset. Our experiment points out new directions for QA problem and our method could be widely used for protein structure prediction problem. The software is freely available at GitHub: https://github.com/caorenzhi/AngularQA

## Introduction

Protein folding prediction proves to be a major hurdle in modern biology. While the rate at which genomes can be sequenced has grown rapidly with the advent of automated systems, protein structures have still been limited to expensive, experimental observation through Nuclear Magnetic Resonance or X-ray crystallography (Jacobson and Sali 2004). While great progress has been made in computational prediction methods with the help of machine learning techniques (Manavalan et al. 2017; Lai et al. 2017; Dao et al. 2017; Qiu et al. 2017; Peterson et al. 2017; Shin, Christoffer, and Kihara 2017), a long journey still remains.

As biology and medicine progresses, the need for a method of reliably and efficiently predicting tertiary protein structures becomes more apparent. Perhaps the most promising use of ab initio folding prediction is in the use of functional prediction and drug discovery (Jacobson and Sali 2004).

The prediction process can be divided into two parts, first generating a model of the target based on its sequence, and then determining how accurate the generated model is (Li, Cao, and Cheng 2015).

Protein structure prediction is usually categorized as being either template-based modeling such as DeepFold (Li, Cao, and Cheng 2015; Liu et al. 2017), FALCON (Wang et al. 2015), MTMG (Li and Cheng 2016), and I-TASSER (Roy, Kucukural, and Zhang 2010); or template-free (ab initio) modeling such as QUARK (Roy, Kucukural, and Zhang 2010; Xu and Zhang 2012) and UniCon3D (Bhattacharya, Cao, and Cheng 2016).

Especially with ab initio modeling, the challenge is then to assess and rank the generated models to help improve prediction, and to know when an acceptable model has been generated (Li, Cao, and Cheng 2015).

Furthermore, these QA methods can be subdivided into two distinct approaches. The first is consensus, which considers many generated models and seeks out patterns to predict which one is the best. This has been shown to work very well with a good dataset generated by multiple different methods, but can be a bad predictor with a poor data-set or small pool and is computationally costly, often requiring O(*n* ^2^) computations where *n* is the number of models (Cao et al. 2016)(Cao, Wang, and Cheng 2014). Such methods include Pcons5(Wallner and Elofsson 2005) and ModFOLDClust2 (McGuffin, Buenavista, and Roche 2013).

The second form of QA, the focus of our research, is single protein assessment. Rather than attempting to score a protein relative to others, the goal is to consider it alone and predict how close it is to the unknown, native structure. Such methods of single-protein assessment include DeepQA which makes use of deep belief networks (Cao et al. 2016), ProQ3 which combines the results from Rosetta energy functions using full-atom and centroid models and the ProQ2 SVM (Uziela et al. 2016), SVMQA which uses a support vector machine to process 19 extracted features (Manavalan and Lee 2017), RFMQA (Manavalan, Lee, and Lee 2014) which ranks model based on random forest, and GMQ (Shin et al. 2017)evaluates local quality based on spatially neighboring residues using a graph representation.

What makes AngularQA especially interesting, is it bypasses most of the costs associated with using two or three dimensional data, reducing a complicated protein with thousands of atoms into a sequence of amino acids, angles, secondary structure, and proximity counts. In addition, only observable features are used in our method further cutting setup costs. The ability of AngularQA is that of its features, without reliance on other, unreliable predictions. This combined with its new methods and use of new features makes it not only a novel approach to single protein quality assessment, but also means it should be of high value to composite QA approaches.

## Method and Implementation

The core of our machine learning model, is a LSTM network which processes each residue and its associated information as a time-step before finally generating an estimated score of the model accuracy. To the best of our knowledge, we are the first to use a recurrent neural network in protein QA.

### Initial Data Preparation

All data used in training comes from 3DRobot decoys generated by The Yang Zhang Lab (Deng, Jia, and Zhang 2016) and from CASP 9, 10, and 11 (Moult et al. 1995). These have 92,535, 36,083, 15,901, and 14,193 models respectively from which we draw for training. Validation occurs on the CASP12, of which we use 6,790 models across 40 targets (Moult et al. 1995). We begin by filtering all the models. During this process we verify the residue sequences in the predicted structures line up correctly with the native structure, and throw out any predicted models with gaps in the center. We also trim the beginnings and ends of the model to line up with the native. In addition, We throw out any models for which we do not have the native structure. After filtering, we are left with a total of 128,439 models with 121,875 training models and 6564 validation models.

We then calculate the GDT scores using the Local-Global Alignment program which superimposes two protein models and assesses the similarity between them (Zemla 2003). Presently, we do not use local alignment scores in training or validation of our method and focus on global quality prediction. All scores are calculated by comparing a given model to the native structure.

Next, we calculate the angles and bond lengths along the backbone and side-chain as was described by UniCon3D; the result is a sequence of angle and bond length information provided for each residue following along the carbon backbone (Bhattacharya, Cao, and Cheng 2016). This representation is central to our method, and has not been extensively studied for its usefulness in QA applications. Because the length of the angle sequence is the same as the length of the protein sequence, the angle information fits well with a recurrent neural network which can work well with inputs of varying lengths (Hochreiter and Schmidhuber 1997). The proximity counts are also calculated at this time by counting the number of C atoms within α a set radius of each residue’s Catom. We perform this calculation for all radii in the discrete α range [5Å, 15Å].

Finally the secondary structures are calculated for all the models using the DSSP program which does not predict the structure, but interprets what is displayed in the predicted model [DSSP]. From its output, we extract only the secondary structure, one of Alpha Helix, Beta Bridge, Strand, Helix-3, Helix-5, Turn, Bend, or if no structure is assigned, Random Coil. The result, is for each residue in the sequence, there is assigned the secondary structure it forms.

### Run-Time Data Preparation

With the great variance allowed for by the PDB format, sometimes one of the initial steps fails for a certain model or group of models, we have added error checking and handling when loading the data to catch inconsistencies and cases where one or more parts may be outright missing. These cases impact less than 1% of models, so we have chosen to identify them, and ignore them. The CASP12 dataset has proven a convenient exception to these challenges, and from our experience, has no such issues.

Before we use the data, we normalize all the angle values we calculate to be in the range [0, 1] and trim the first and last residues from each model. The values at both ends we found to have extreme values in many cases which are not related to the sequence.

Additionally, because proteins have different numbers of residues, we pad the data to make the lengths consistent for training; the model itself later masks zero values to counteract this. We choose to pad at a length of 500 as most proteins in our dataset are shorter. Of the 688 targets in our dataset (including CASP12), they have a distribution of N(180.0, 119.6) with only 13 longer than 500 (This average is based on the length of the observed structures which are sometimes missing a few residues at the front or end.). Those few which are longer drop the last residues. A notable benefit to our representations, is their small space requirement; when loaded, all our datasets combined, roughly 160,000 models, use less than 4GB of memory even when padded to a length of 500 residues.

### Features

Each time-step includes information about that residue. The amino acid is considered one of the most fundamental and is included with all tests.

The core features we use are the angles between residues. To verify this new representation is of value, we calculated the correlation between the different angles---Tau, Theta, Phi, and Delta---and the related GDT score finding weak correlations for both Cα angles (r_τ_=0.373, r_⎕_=0.427) and ατ lesser correlations for both side-chain angles (r_⎕_=0.187, r_δ_=0.299); these results indicate the δ angles within and between residues could be a good feature for assessing predicted structures. DSSP determines the secondary structures at each of the residues based on their 3D form and is then fed in along side the amino acid in addition to the results from Proxcalc, an in-house program to calculate and count the residues within a given radius (Joosten et al. 2011).

We have tried different combinations of these, and for now have settled on using the amino acid, theta, tau, the secondary structure, protein properties, and proximity counts for radii of 8Å and 12Å.

### LSTM Network

The network begins by masking the zero values of padded proteins. Followed after, the data for each time-step is separated into three parts: the amino acid, the secondary structure, and physical protein properties (hydrophobic, polar, or charged) and the angles and bond lengths. The amino acid and secondary structure are then converted to dense-vector encodings, mapping the value to n-dimensional space---in both cases we use four dimensions.

With the vectorized values, they are then reunited with the other data, forming the true LSTM input for each time-step. The LSTM layers vary in breadth and depth, but a common configuration we test which achieves highly, is a [64, 32, 10] arrangement. The first two layers return a sequence of values, one for each time step, while the final layer outputs a single value at the end of the residue series. Each LSTM cells uses a hyperbolic tangent activation with a hard sigmoid recurrent activation.

The output of the final layer then gets run through a single layer of perceptrons with sigmoid activation and learned weights before being converted to a single value in the range (0, 1). The network is trained using RMSprop with a learning rate of 0.0001. Stochastic gradient descent and Nadam were also tried, but neither proved as stable or effective.

## Results and Discussion

Thus far, we have achieved an overall correlation of 0.684 on CASP12 with an average loss of 0.122. These results were found using three LSTM layers in the configuration [128, 64, 32] which was trained for less than 154 epochs on all models from 3DRobot and CASP 9, 10, and 11.

**Table 1** and **Table 2** demonstrate the Pearson Correlation and loss before and after trimming Stage 1 and Stage 2 datasets for CASP12. We found a surprising difference between running on the CASP12 data which had been trimmed to the native structure (If the native was missing part of the sequence at the beginning or end, it was removed for all trimmed tests.) versus our results with the raw predictions. Of course, using the trimmed data requires more information than is available in real-world and thus represents no more than an interesting comparison. In addition, we compare performance of our method AngularQA with few selected top performing methods from CASP12. **Table 3** describes the average per-target Pearson Correlation and loss for our method AngularQA and four selected top performing single-model QA methods. We could see that ProQ3 method achieves the best performance among all methods. It is not very surprising that ProQ3 performs better than our method, because our method only uses the information from the model, such as angle information. We don’t even use the secondary structure prediction from the sequence, which is usually a very useful feature for model quality assessment. At the same time, we also find out that AngularQA performs better the other two methods Wang1 and QMEAN. The similar pattern is found in Stage 2 datasets, which is shown in **Table 4**. Overall, based on our experiment, our AngularQA method demonstrates a great potential of using angle information for model quality assessment, and our method could be even furtherly used as a feature for top performing QA method and improve the accuracy of these QA method.

**Table 1.**
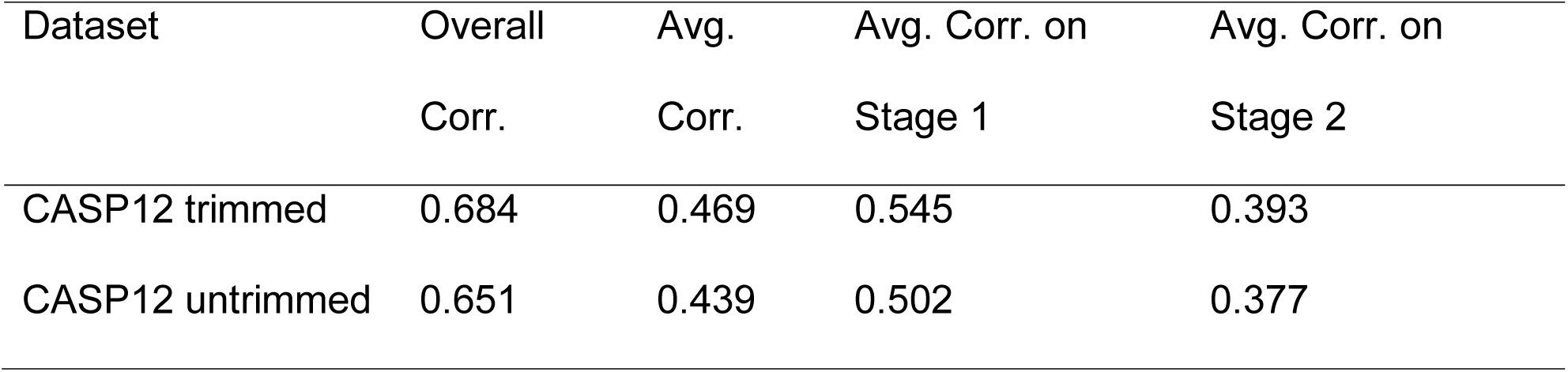
Comparison of correlations between datasets on Stage 1 and Stage 2 of CASP12. Overall correlation uses all data points while the average correlation is the mean of correlation scores.

**Table 2.** Comparison of loss scores between datasets on Stage 1 and Stage 2 of CASP12.

| Dataset | Ave. Loss | Loss on Stage 1 | Loss on Stage 2 |
| --- | --- | --- | --- |
| CASP12 trimmed | 0.122 | 0.116 | 0.128 |
| CASP12 untrimmed | 0.138 | 0.148 | 0.128 |

**Table 3.**
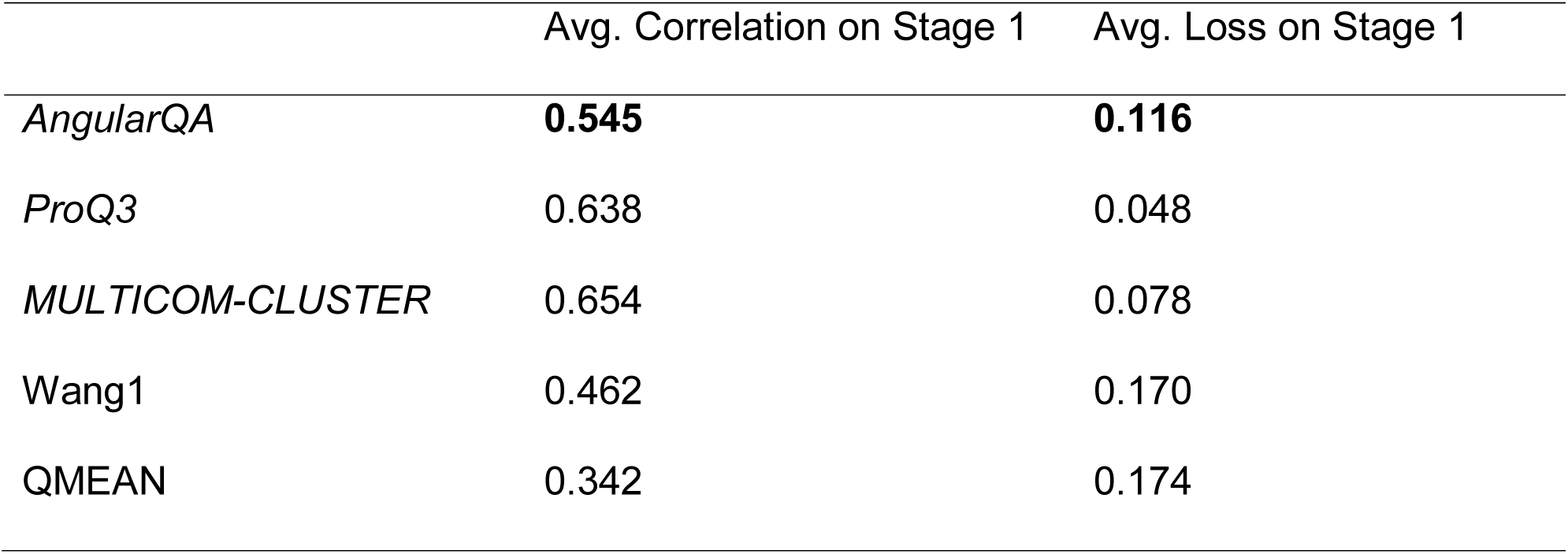
The performance of the global quality predictions of our AngularQA method and four selected CSP12 methods in terms of average correlation, and average loss of top 1 models ranked by each method, evaluated on CASP12 Stage 1 targets.

**Table 4.** The performance of the global quality predictions of our AngularQA method and four selected CSP12 methods in terms of average correlation, and average loss of top 1 models ranked by each method, evaluated on CASP12 Stage 2 targets.

|  | Avg. Correlation on Stage 2 | Avg. Loss on Stage 2 |
| --- | --- | --- |
| *AngularQA* | 0.393 | 0.128 |
| *ProQ3* | **0.616** | **0.068** |
| *MULTICOM-CLUSTER* | 0.578 | 0.100 |
| Wang1 | 0.256 | 0.144 |
| QMEAN | 0.292 | 0.125 |

## Conclusions

Our results show great promise for the use of both angle information in QA, as well as recurrent neural networks. The angle correlations we calculated show they can be a useful metric in protein quality assessment. Our work demonstrates these values are generalizable between models of the same target, and more importantly to new, unseen targets.

Interestingly, before we added the secondary structure information, the overall correlations between the true and predicted scores for all attempted networks were below 0.3 before adding the secondary structure information and were often closer to 0.15. This indicates the secondary structure helps the model assess the validity of angles and determine if the overall is coherent. While we have tested different combinations and ablations of both features and layer setups, much work remains to optimize the system as a whole. The thing we found to have had the largest impact beyond the features and data, was the learning rate which we ended up reducing by a two factors of ten. Before doing so, we found the models trained in very few epochs, somewhere between 10 and 40 in most cases. Since reducing the learning rate we have found the network to perform less well on training data, but to perform much better on testing data, and begin overfitting after a couple hundred epochs.

To help reduce the rate of drift in training, we wanted to increase the number of models we were using for validation, however, when we tried blindly splitting the training set for validation, we found the model scored very highly while training it, but when we went to test it against only CASP12, its performance was far worse than without the extra testing data points. This indicates it was able to recognize similar structures to what it was trained on and adequately assess them. To reduce this issue, we could make sure data splits take entire targets at a time rather than individual models preventing it from training on models from some of the same targets it will later be validated on.

In future, new features could be added to further help the system, such as the physical properties of the amino acids or contact information. We have also considered changing the way it runs, possibly adding a bidirectional LSTM system to consider the sequence from both ends. Overall, there are many possibilities left to try, and our work shows QA based on angles and using recurrent neural networks has great promise.

**Figure 1.**
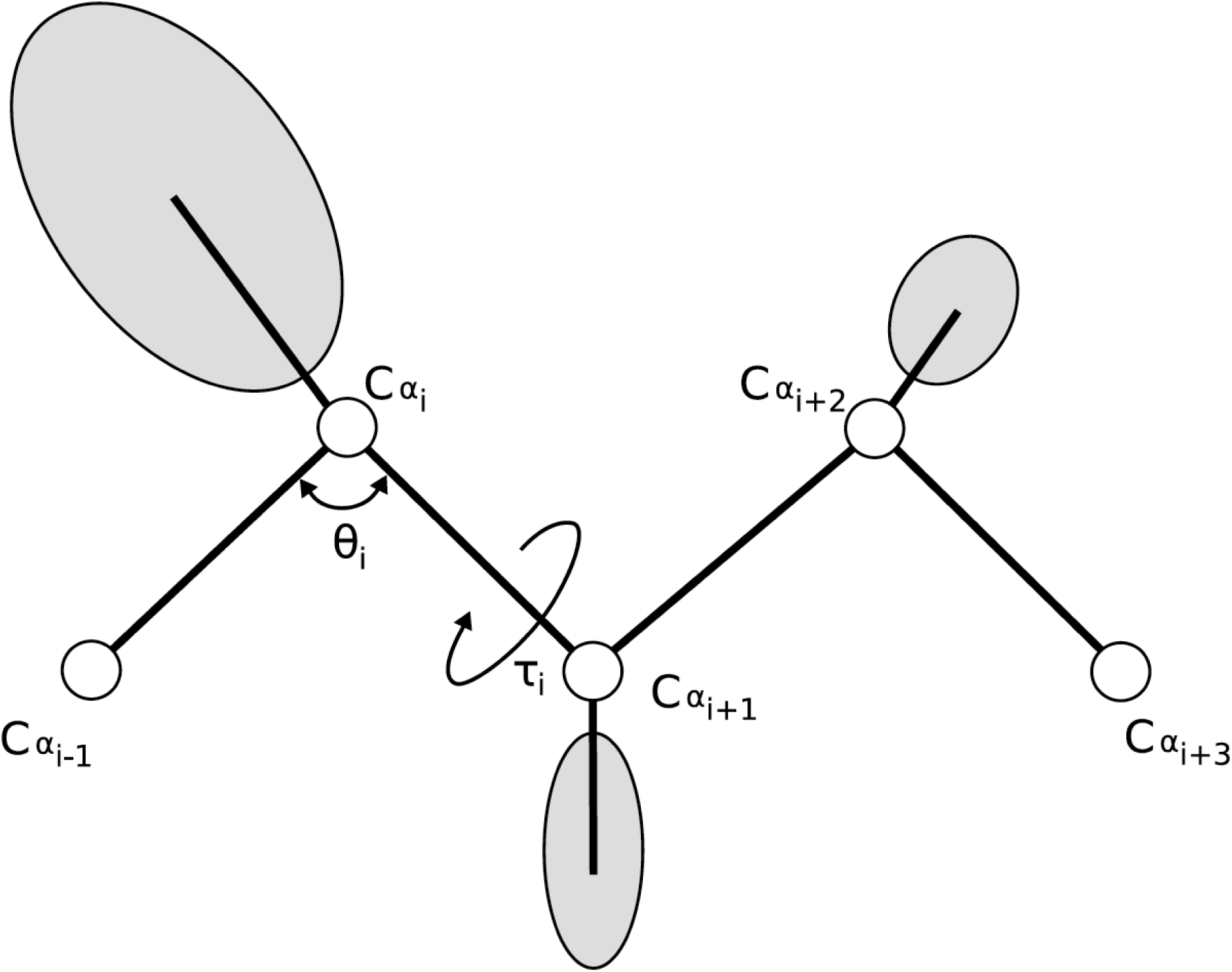
Diagram of the LSTM network components and data flow.

**Figure 2.**
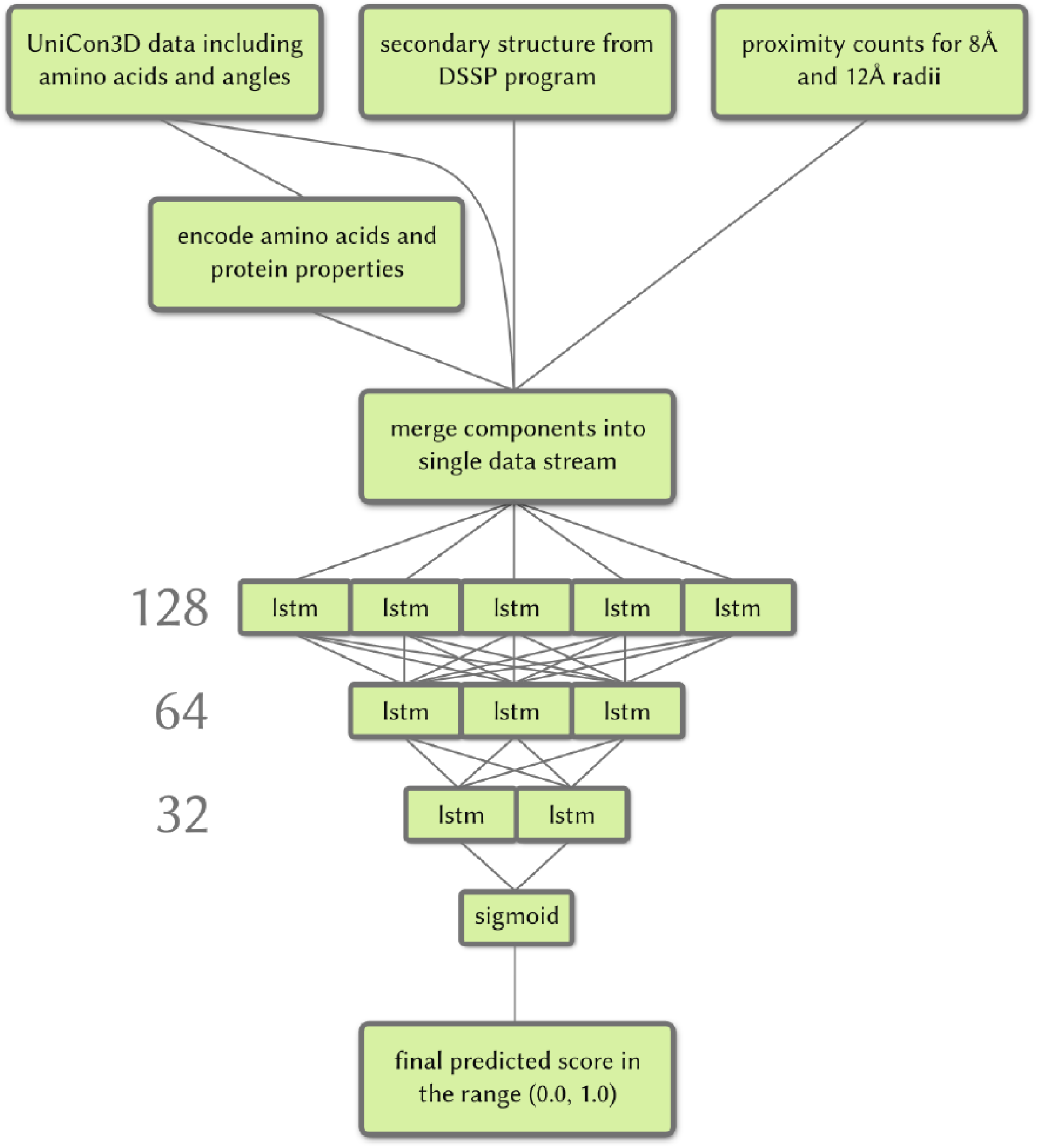
Representation of angles and bond-lengths. Each Cα is interpreted as an individual time-step for the network.

## Authors’ contributions

RC conceived the project. MC, RC designed the project. MC and RC implemented and tested the tool. MC, DS, MS, RC wrote the manuscript. All the authors read and approved the manuscript.

## Acknowledgements

The authors would like to thank support from students and faculties at Pacific Lutheran University. Also, we thank the anonymous referees for their valuable comments and helpful suggestions. This work is supported by Natural Sciences Undergraduate Research Program at Pacific Lutheran University, Karen Hille Phillips Regency Advancement Award to Prof. Cao, and the Division of Natural Science at Pacific Lutheran University.

